# A novel growth and isolation medium for exoelectrogenic bacteria

**DOI:** 10.1101/2021.11.17.469056

**Authors:** Zumaira Nazeer, Eustace Y. Fernando

**Affiliations:** Faculty of Applied Sciences, Department of Biological Sciences, Rajarata University, Mihintale, 50300, Sri Lanka

**Keywords:** Growth medium, selective medium, exoelectrogenic bacteria, microbial fuel cell

## Abstract

A microbiological isolation and growth medium that can effectively discriminate electrochemically active exoelectrogenic bacteria from other non-exoelectrogenic bacteria, is currently unavailable. In this study, we developed a novel chromogenic growth and isolation solid medium based on MnO_2_ that can selectively allow the growth of exoelectrogenic bacteria and change the medium colour in the process. Known exoelectrogenic bacteria such as *Shewanella oneidensis* MR1 and other such bacteria from functional microbial fuel cell (MFC) anodes were capable of growing and changing colour in the novel growth medium. On the contrary, non-exoelectrogenic bacteria such as *Escherichia coli* ATCC 25922 were incapable of growing and inducing a colour change in the novel medium. Further biochemical characterisation of these isolated exoelectrogenic bacteria by Raman micro-spectroscopy demonstrated that these bacteria over express cytochrome proteins that are vital in extracellular electron transfer events. This medium is a convenient method to isolate exoelectrogenic bacteria from complex environmental samples.

## 1. Introduction

Exoelectrogenic bacteria are characterized by their special capability of transferring metabolic electrons beyond the cellular boundary to electron acceptors that reside outside (Jiang et al., 2016 and Logan., 2009). Most often this extracellular electron transfer (EET) takes place onto insoluble reduced electron acceptors such as naturally available metal oxides (Iron and Manganese oxides). Exoelectrogenic bacteria are the driving force behind bio-electrochemical devices such as microbial fuel cells (MFC), microbial electrolysis cells (MES) and microbial electro-synthesis systems (Wang et al., 2019). At present, many exoelectrogenic bacteria are known to exist in monoculture. The best characterised and the most widely used include species such as *Geobacter sulfurreducens, Geobacter metallireducens* and *Shewanella oneidensis* among many other bacterial species (Cheng and Call., 2021). These exoelectrogenic bacteria often form complex communities in their natural habitats. Therefore, isolation and characterization of exoelectrogenic bacteria in monoculture was shown to be difficult (Becerril-Varela et al., 2021). Many previous studies have utilized environmental mixed microbial consortia in their work with little or no characterization of the underlying exoelectrogenic bacterial community. Such sources of exoelectrogens forming complex communities include lake sediments, marine sediments, anaerobic digester sludge and many other mixed microbial consortia from both natural and engineered habitats (Fernando et al., 2021). Engineered habitats harbouring exoelectrogens are mainly MFCs and microbial electrolysis cells (MECs).

At present, no custom-designed growth and isolation medium for isolation of exoelectrogenic bacteria exists. This study was specifically designed to address that gap and to produce an effective growth and isolation medium for selecting exoelectrogenic bacteria. Such a medium should be capable of supporting the specific type of metabolism based on EET events and should be effective at differentiating the EET-based metabolism from all other metabolic activities by the microbial consortium.

The objective of this study was to design and characterize a chromogenic isolation and growth medium for exoelectrogenic bacteria. An insoluble electron acceptor such as MnO_2_ that exists as a black mineral precipitate when oxidised (Mn^4+^) and a colourless, more soluble state when reduced (Mn^2+^) will act as an ideal candidate for designing such a chromogenic medium that is capable of isolating exoelectrogenic bacteria. Therefore, this study utilized a MnO_2_ (Mn^4+^) – based chromogenic minimal growth medium to effectively grow and isolate exoelectrogenic bacteria from a complex environmental bacterial consortium.

## 2. Materials and methods

### 2.1. Lake sediment sample collection

The tropical lake sediment mixed bacterial consortium was collected from the ancient man-made lake Mihintale, Sri Lanka, as previously described (Dissanayake et al., 2021) (located 110 m above sea level; 8.3640°N, 80.5085°E). Approximately 10 g of sediments from about 10 cm below the lake sediment surface were collected in a sterile cap-secured sample collecting tube and transported to the laboratory at 4°C. At the time of sampling, the lake sediment indicated moderate anaerobic activity and some pyrite mineral deposition. The samples were kept airtight and opened only when inoculating MFCs.

### 2.2. Chemicals, reagents and bacterial strains

All chemicals used in media formulations were of analytical grade and were procured from Sigma Aldrich, UK. The reference electrochemically active bacterium *Shewanella oneidensis* strain MR-1 was procured from the NCIMB culture collection, UK. The reference non-electrochemically active bacterium *Escherichia coli* ATCC 25922 was procured from the culture collection ATCC, USA.

### 2.3. Inoculation of lake sediment bacteria and operation of MFCs

Exoelectrogenic bacteria were initially selected in anode compartments of microbial fuel cells (MFCs) that operated continuously for more than four months. MFC anodes were inoculated with approximately 10 cm^3^ of tropical lake sediment. Anode chambers of MFCs were filled with 250 mL of a semi-defined minimal media with Glucose as the main carbon source. The semi-defined anode medium was formulated with Glucose 2g /L, NH_4_Cl 0.46 g/L, yeast extract 0.1 g/L, peptone 0.5 g/L, K_2_HPO_4_ 5.05 g/L, KH_2_PO_4_ 2.84 g/L (pH −7.1). The MFCs containing two compartments were designed as shown in the schematic (Figure-01). They were constructed using cast acrylic sheet, with appropriate measurements so that the working volume of each compartment was approximately 250cm^3^. The two compartments were separated by pre-treated Nafion™ 212 proton exchange membrane as described earlier by (Logan et al., 2006). Phosphate buffer solution of 50mM strength, pH 7.1 served as the catholyte in the cathode compartment. The cathode compartment was continuously aerated at a constant air flow rate of 100 mL/min using an air pump.

**Figure - 1:**
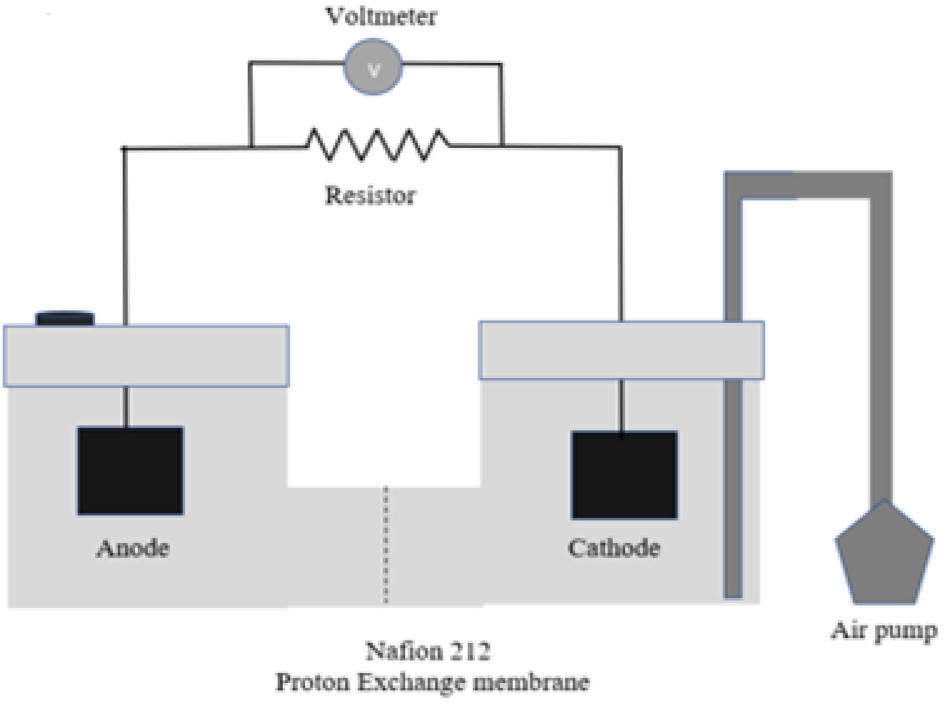
a schematic diagram of the two-chamber MFC system used in this study for obtaining electrochemically active anode bacteria.

A 5cm × 5cm conductive carbon cloth was used as electrodes in both cathode and anode. Electrodes were connected externally by insulated copper wire, with resistors of 680 Ω in between. A PicoLog™ 1012 programmable data logging voltmeter (Pico Technologies, UK) was attached in parallel to the resistor. Voltage data across the 100 Ω external resistances was automatically logged and stored at an interval of 10 minutes for the duration of MFC experiments.

Platinum nano-powder (Sigma Aldrich, UK) was coated on the cathode surface as the MFC cathode catalyst at a loading of 0.5 mg/cm^2^ using Nafion™ conductive perfluorinated resin solution as described earlier (Fernando et al., 2013). Performance characterization of MFCs using polarization curves and power-current plots was conducted as described earlier (Fernando et al., 2016). MFCs were operated in semi-continuous mode with substrate feeding for more than two months before obtaining samples of exoelectrogenic bacteria for isolation experiments.

### 2.4. Novel chromogenic growth and isolation medium for exoelectrogenic bacteria

The novel chromogenic medium for isolation and growth of exoelectrogenic bacteria composed of the following components (g/L); Glucose - 2, NH_4_Cl - 0.46, yeast extract - 0.1, peptone - 0.5, K_2_HPO_4_ - 5.05, KH_2_PO_4_ - 2.84, MNO_2-_ 1.5, cysteine - 0.025 and agar - 15. Cysteine was used in the growth medium to scavenge residual oxygen during incubations. This medium was designed to isolate microbes which can utilize MNO_2_ as the final electron acceptor. Solid MnO_2_ fine particles were suspended in molten agar medium before it was poured in Petri dishes. This produced a uniform jet-black colour in the novel medium when it was fully set, prior to inoculation of bacteria (Figure 3A). The MFC anode biofilm bacteria from functioning MFCs were introduced into the solid novel minimal medium by streak plating. The plates were incubated under an anaerobic atmosphere in an anaerobic jar (Oxoid, UK). The anaerobic atmosphere inside the sealed jar was generated by the use of Gas-Pak™ anaerobic sachets. *Escherichia coli* type strain ATCC 25922 was grown in the same medium under the same conditions as the reference bacterial culture.

### 2.5. Raman micro-spectroscopic characterization of MFC anode biofilms and isolated bacteria

Biofilm samples from MFC anodes and isolated colonies from novel growth medium were carefully scraped using a sterile sharp scalpel and were immediately preserved in ice-cold 50% (V/V) ethanol/PBS (50 mM, pH – 7) for Raman analysis. Preserved biofilm samples were mounted and analysed on CaF_2_ Raman windows (Crysran, UK), as described previously (Fernando et al., 2019). Briefly, Raman microspectroscopy was conducted using a Reneshaw™ InVia (UK) Raman microscope and the Raman excitation laser beam had a wavelength of 514 nm. The diffraction grating of the Raman instrument used during all analyses was a 1200 grooves/mm. The incident Raman laser beam was attenuated down to 5% of the total power using a neutral density (ND) filter. The excitation laser of the Raman system was focused onto the biofilm sample using a 40 × air objective with a numerical aperture of 1.0. The return Raman signal was collected using the same objective and the Raman spectra were recorded on a CCD detector cooled to - 70°C. Raman spectra from the biofilm samples were collected within the spectral range of 200 cm^−1^ to 1700 cm^−1^. The Raman system was calibrated to the first-order scattering signal of a standard silicon wafer at 520.7 cm^−1^. All spectra were recorded using the in-built Wire 5.0 software of the Raman system (Reneshaw Instruments, UK). *E. coli* ATCC 25922 cells grown in nutrient broth was also used for reference in Raman analysis.

Raman mapping experiments were conducted in all three dimensions of the biofilm sample (X, Y and Z planes) by assigning a raster pattern of 0.5μm step size spot measurements over a total mapping area of 16μm^2^. The sampling interval in the depth (Z) plane was 0.3 μm. This allowed for a Raman pseudo-colour 3D map of exoelectrogenic bacteria to be generated.

### 2.6. Molecular microbial characterisation and phylogeny of anode exoelectrogenic bacteria

Isolated exoelectrogenic bacteria on the novel chromogenic medium were subjected to molecular characterization by using the 16s ribosomal RNA (16s rRNA) marker gene, as described earlier (Dissanayake et al., 2021). Briefly, the total genomic DNA from single colonies of isolated bacteria from the custom designed selective medium described in the section 2.3 was conducted using a Wizard™ genomic DNA extraction kit (Promega Corporation, USA), as per the manufacturer instructions. The 16s rRNA gene was amplified from the extracted genomic DNA using the primer pair 27F (5’ - AGA GTT TGA TCM TGG CTC AG - 3’) and 907R (5’-CCG TCA ATT CCT TTR AGT TT– 3). The PCR master mix (Promega, USA) was used as per the manufacturer instructions to amplify the 16s rRNA marker gene, in a PCR thermocycler (Thermo-Fisher, UK) under the following set of conditions; initial denaturation at 95°C for 4 min, followed by 30 cycles of 95°C for 0.5 min, 58°C for 1 min, 72°C for 0.5 min, and finally at 72°C for 7 min. PCR products were confirmed and verified on 1% agarose gels before being sequenced using the Sanger DNA sequencing technology at the Department of Molecular Biology, University of Peradeniya, Sri Lanka. Nucleotide sequences obtained from bidirectional sequences were checked for chimeras using the software DECIPHER and the consensus sequences from bidirectional raw reads were assembled using the software BioEdit version 7.0.5.3. Fully assembled and quality checked consensus sequences were compared with the 16s rRNA gene entries of the databases NCBI GenBank 16S rRNA nucleotide sequence repository and Ribosomal Database Project (RDP) using NCBI BLAST nucleotide search tool (http://blast.ncbi.nlm.nih.gov/Blast). Nucleotide sequences obtained from the isolated exoelectrogenic bacteria were deposited in the NCBI GenBank 16s rRNA sequences repository, under the accession numbers OK083658-OK083662. Phylogenetic analysis of the sequences was conducted as described earlier (Dissanayake et al., 2021 and Fernando et al., 2013), using Mega 7.0.026 molecular evolutionary genetics analysis software with maximum likelihood method (1000 bootstrap replicates).

## 3. Results and discussion

### 3.1. The operation and performance of microbial fuel cells

The initial acclimation and selection of exoelectrogenic bacteria from tropical lake sediment was done in two-chambered microbial fuel cells operating in semi-continuous operation. The MFCs exhibited a very repeatable and a stable operation over long periods of time. They exhibited an open-circuit voltage of approximately 500 mV. During stable operation, the average maximum power density (P_max_) obtainable was approximately 180 mWm^−2^ and the maximum current density (J_max_) was approximately 1058 mAm^−2^ (Figure - 2B). The MFCs had a stable closed-circuit voltage across an external resistance of 100Ω over long periods of semi-continuous MFC operation (Figure - 1). This is indicative of stable electrochemical performance of the MFC system. It also suggests that the microbial community of the anode compartment of the MFC was well acclimated and stable during the MFC operation. The internal resistance (R_int_) of the MFC systems used was found to be approximately 90Ω.

**Figure-2:**
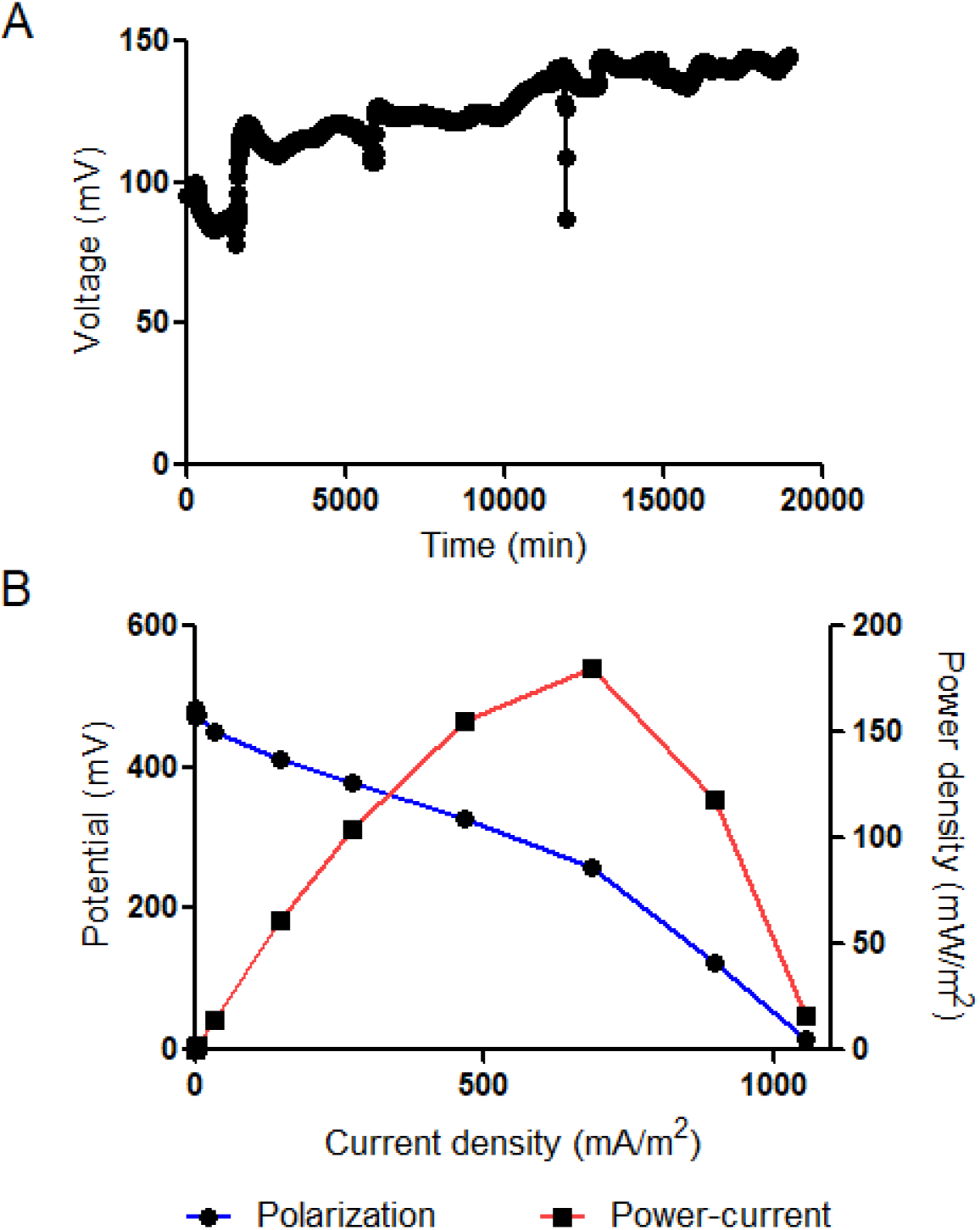
(A) Voltage output of the MFCs (average of 2 MFCs) used in this study during their operation for a period of approximately 13 days (voltage interruptions are due to opening of the circuits for substrate feeding events) (B) polarization curve and the power-current plot of the MFC systems indicating a good electrochemical performance for a two-chamber MFC system (P_max_ = 180 mWm^−2^ and J_max_ = 1080 mAm^−2^).

High current and power densities together with a stable and a sustainable closed-circuit voltage obtained from the MFC system demonstrated that the resident microbial community was well adapted to conducting extracellular electron transfer reactions onto the anode electrode of the MFC.

### 3.2. Isolation of exoelectrogens on novel growth medium

Culture-dependant isolation of electrochemically-active single bacteria was done using a custom designed minimal growth medium that changed colour based-on extracellular electron transfer metabolism. The novel growth medium contained glucose as the sole source of carbon and energy and also provided the reducing equivalents for the extracellular electron transfer reactions. All other nutrients such as nitrogen, sulphur and phosphorus were provided in the novel growth medium as defined components in known quantities. All other electron soluble acceptors such as oxygen, sulphate or nitrate ions were absent in the growth medium. The only electron acceptor provided in the novel growth medium was the insoluble metal oxide electron acceptor MnO_2_. Black coloured manganese dioxide (Manganese (IV) oxide), upon accepting metabolic electrons from substrate oxidation will be converted into colourless and more soluble Mn^2+^ ions as given in expression-1.

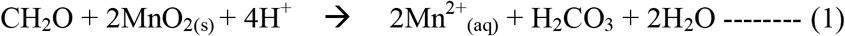

This clear colour change taking place due to MnO2 accepting metabolic electrons from electrochemically active bacteria conducting extracellular electron transfer reactions offers a convenient way to grow and select for exoelectrogenic bacteria under laboratory conditions. Due to the absence of all other electron acceptors such as oxygen (anaerobic atmosphere) and nitrate and sulphate (not provided in the minimal medium), the sole electron acceptor available for any microorganism capable of growing in this novel medium should be capable of utilizing insoluble MnO_2_ as the terminal electron acceptor. This also implies that any microorganism demonstrating growth on this medium must be an exoelectrogenic bacterium due to the fact that metabolic electrons should be transferred across the cellular boundary and into the MnO_2_ granules located in the cellular exterior. Bacteria growing on the novel defined medium were capable of fully or partially decolourizing the black growth medium, depending on their growth rate on the solid medium (Figure - 3). The initial colour of the medium prior to inoculation was black (Figure-3A). When known electrochemically active bacteria such as *S. oneidensis* MR1 were grown in this novel medium, they were able to grow and made the growth medium completely colourless at the end of the growth period of 72 hours (Figure-3B). Bacteria that were selected in MFC anodes for extended operational periods were effectively isolated using this novel growth medium. The isolates grew well on the solid medium and were capable of completely decolourising the black colour in the process (Figure - 3C). Bacteria that were known to possess no extracellular electron transfer capabilities such as the control strain *E. coli* ATCC 25922 was incapable of utilizing and growing in this novel growth medium containing MnO_2_ as the sole electron acceptor (Figure - 3D). Growth of exoelectrogenic bacteria and non exoelectrogenic bacteria on this novel medium is clearly distinguishable by the absence of growth and colour change for non-exoelectrogens (Figure - 3E).

**Figure-3:**
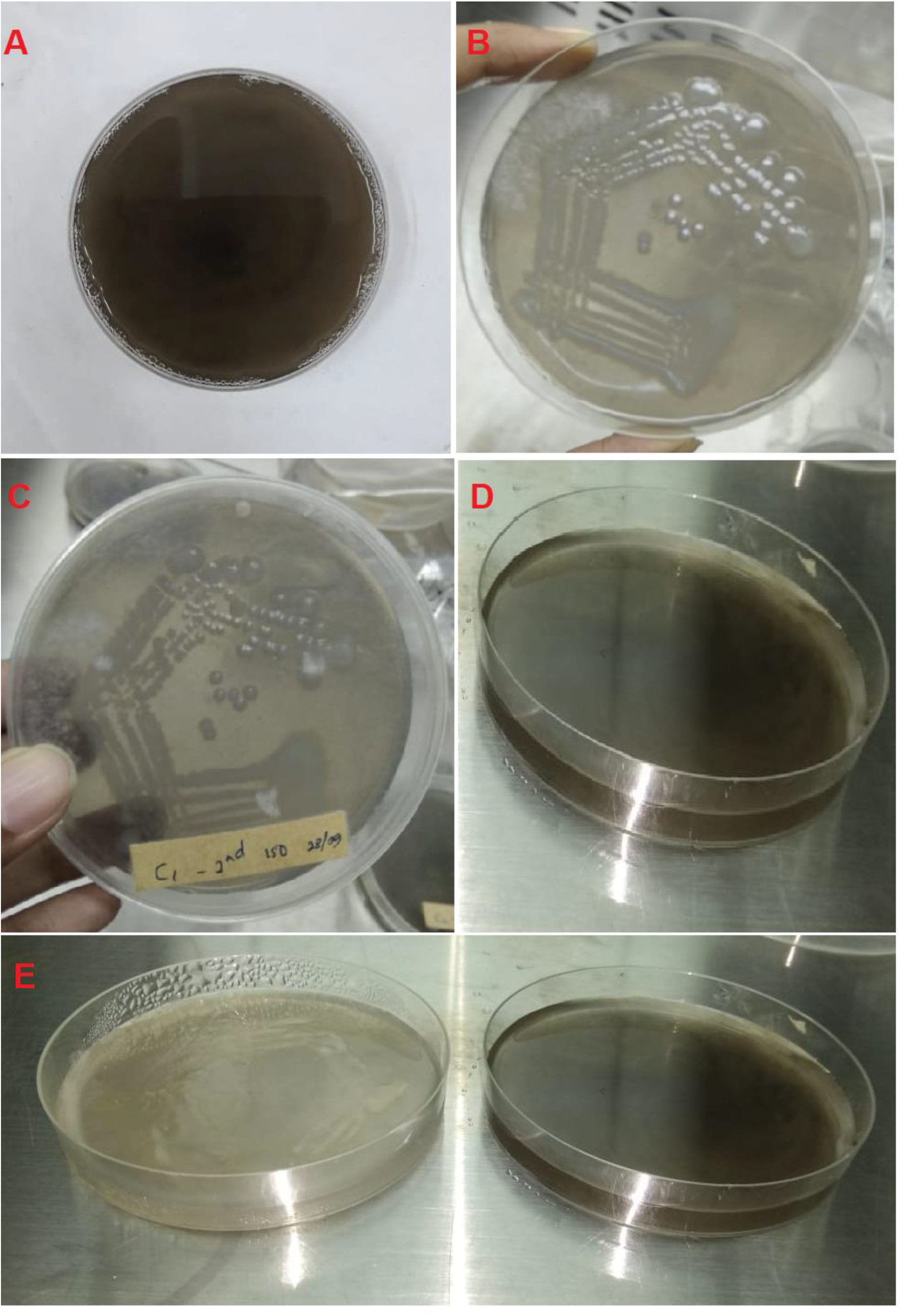
(A) Un-inoculated blank medium indicating the clear black colour of MnO_2_ (B) electrochemically active bacterium *S.oneidensis* MR1 grown on the novel medium, indicating growth and complete colour removal (C) An isolated bacterium from a functioning MFC anode biofilm indicating growth and decolourisation of the novel MnO_2_ medium (later characterized to be *Klebsiella pneumoniae* (D) Absence of growth and decolourisation of the novel MnO_2_ – based medium when *E.coli* ATCC 25922 was inoculated (E) a side-by side comparison of the novel growth medium appearance when utilized by electrochemically active bacteria (*S.oneidensis* MR1) and when it cannot be used by non-exoelectrogens such as *E.coli*.

*all bacterial isolates were grown for 72 hours on the novel MnO_2_ containing solid growth medium

This clearly demonstrates that bacteria incapable of shuttling their metabolic electrons outside their cellular envelope into insoluble terminal electron acceptors in the cellular exterior are incapable of utilizing and growing in this novel growth medium. The non exoelectrogenic reference bacterium *E. coli* ATCC 25922 used in this study isn’t able to shuttle electrons into insoluble electron acceptors situated outside cellular boundaries. Their lack of growth on this novel growth medium can therefore be attributed to the absence of a suitable terminal electron acceptor.

A suitable terminal electron acceptor for such non- exoelectrogenic bacteria would ideally be a soluble chemical species (oxygen, organic acids, nitrate or sulphate ions) that can readily cross cellular boundaries and membranes into the intracellular milieu and accept metabolic electrons resulting from substrate oxidation. MnO_2_ used in this novel growth medium is insoluble, located outside the cellular boundary and therefore, unsuitable for use by any other bacteria other than exoelectrogenic electrochemically active microorganisms. In the event of complete absence of alternative electron acceptors other than MnO_2_, non-exoelectrogenic bacteria will simply fail to utilize the novel solid medium to grow. Electrochemically active exoelectrogenic bacteria on the other hand, readily possess the biochemical and physiological ability to conduct long distance electron transfer to transfer their metabolic electrons outside their intracellular milieu and into an insoluble electron acceptor such as MnO_2_ located outside the cell boundary.

### 3.3. Raman spectroscopic characterization of electrochemically active MFC anode biofilms

Raman micro-spectroscopy is a single-cell microbiology technique that can be employed to show biochemical and prominent molecular features of single cells in their native environment (Berry et al., 2015 and Fernando et al., 2019). The use of Raman micro-spectroscopy for qualitative and quantitative characterization of chemical species such as polyphosphate, polyhydroxyalkanoates and glycogen (Fernando et al., 2019), elemental sulphur (Eichinger et al., 2014) and many other intracellular chemical species in bacteria (Wagner., 2014) has been demonstrated in previous studies.

It has also been demonstrated that cytochrome proteins that are involved in extracellular electron transfer in bacteria produce a very distinct Raman spectroscopic signature in their reduced state (Haider et al., 2010 and Virdis et al., 2014). Excess presence of reduced cytochrome proteins in cell surfaces and intracellular environments can be conveniently detected by the concurrent presence of four very prominent Raman peaks at positions 750 cm^−1^, 1129cm^−1^, 1311 cm^−1^ and 1585 cm^−1^ (Pätzold et al., 2008) The strong concurrent presence of these Raman peaks is indicative of the excessive presence and usage of reduced cytochrome proteins of cells for electron shuttling purposes.

This outcomes of study clearly shows that a strong presence of all four characteristic peaks for reduced cytochrome proteins are identifiable in in Raman spectra of the known electrochemically active bacterium *S.oneidensis* MR1 cells grown in the novel MnO_2_ containing solid medium (Figure 4A, spectrum-I). Similarly, the bacterial cells isolated from functional MFC anodes and later grown in the novel MnO_2_ containing solid growth medium also exhibited the same Raman spectral profile containing the signature peaks at 750 cm^−1^, 1129 cm^−1^, 1311 cm^−1^ and 1585 cm^−1^ that can be directly attributed to the abundant expression of cytochrome proteins (Figure 4A, spectrum-II). These isolated bacterial cells analysed by Raman micro-spectroscopy were later identified by 16s rRNA gene molecular studies to be *Klebsiella pneumoniae* (GenBank accession number - OK083658). By contrast, the known non-exoelectrogenic bacterium *E.coli* produced a Raman spectrum completely devoid of any peaks that can be attributed to the excessive presence of reduced cytochrome proteins (Figure 4A, spectrum-III). Characterization of an entire area of a MFC anode biofilm revealed that some cells of the cell population of anode biofilm over-expressed the cytochrome proteins that are vital in long-distance extracellular electron transfer (Figure 3B). The outcome of this Raman mapping experiment (Figure - 4B) of the MFC anode biofilm clearly demonstrated a heavy presence of the signature Raman peaks for reduced cytochrome proteins. Superimposition of some of these signature peaks (750 cm^−1^ assigned blue and 1585 cm^−1^ assigned green), highlighted the cells in the MFC anode biofilm that overexpress these cytochrome proteins in the Raman pseudocolour map (Figure - 4B and 4C). The cyan areas of the Raman map represent overlap of Green and Blue Raman signal; indicating the presence and localization of the reduced cytochrome proteins used by certain populations of bacteria within the MFC anode biofilm for extracellular electron shuttling events.

**Figure - 4:**
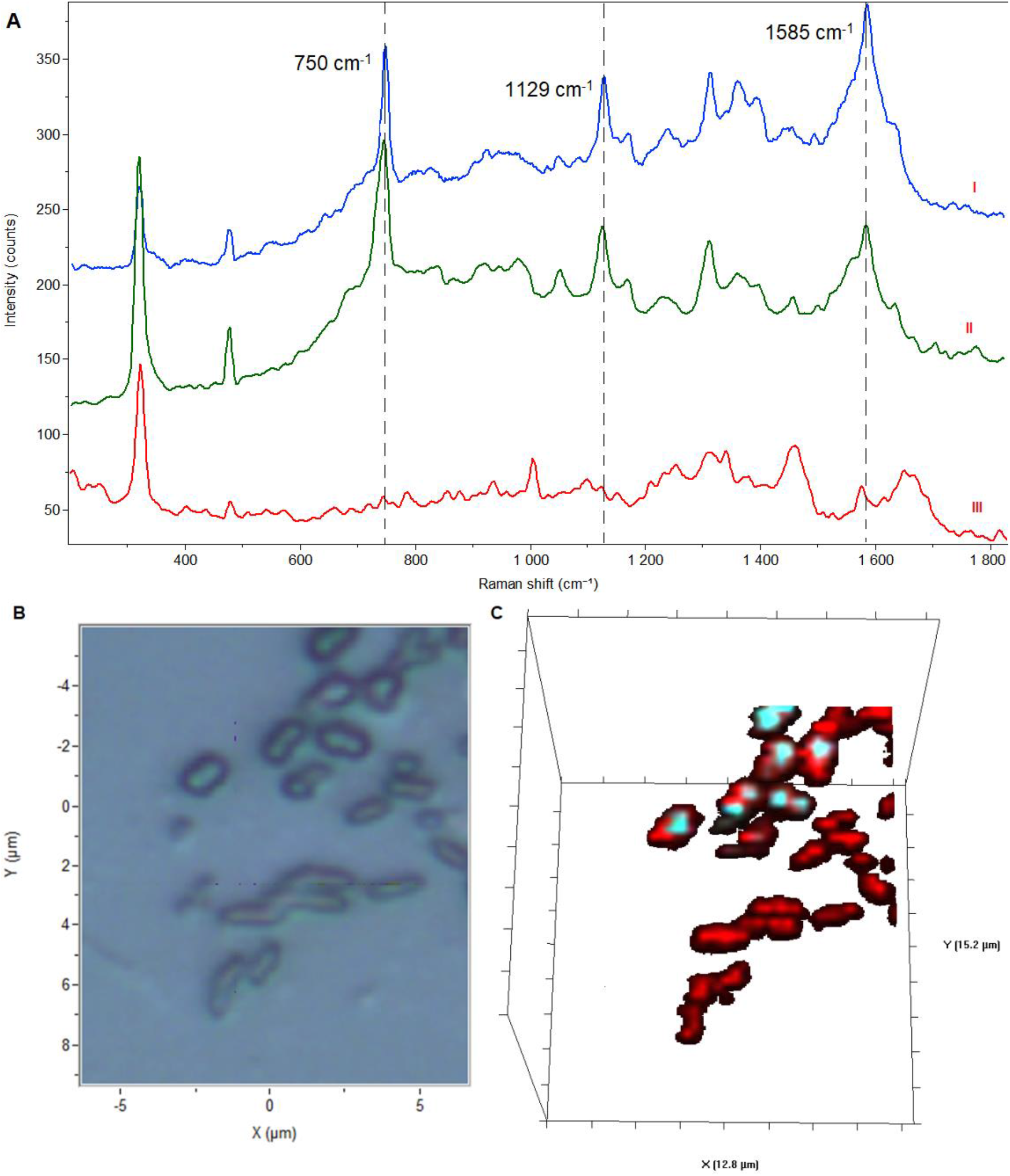
(A) A Raman spectral overlay representing S.oneidensis MR1 cells grown on the novel MnO_2_ containing solid medium (spectrum-I), Raman spectrum from an isolated bacterium from the MFC anode biofilm that later isolated on the novel MnO_2_ containing solid medium (spectrum-II) and the average Raman spectrum from non-exoelectrogenic reference bacterium *E. coli* ATCC 25922 (spectrum-III) (all representative spectra are averages of 10 individual Raman spectra) (B) Brightfield microscopic image of an area of the MFC anode biofilm cells that were analysed by a 3-dimensional (3D) Raman mapping experiment (C) Raman pseudocolour 3D mapping experiment of the area of the MFC anode biofilm shown in the brightfield image. Colour assignment for Raman peaks – Red - 1660 cm^−1^ (stretching vibrations of amide III linkages representing general biomass), Green – 750 cm^−1^ (stretching vibrations of the pyrrole ring of reduced cytochromes), Blue – 1129 cm^−1^ (C-N stretching vibrations of reduced cytochrome proteins).

**Figure - 5:**
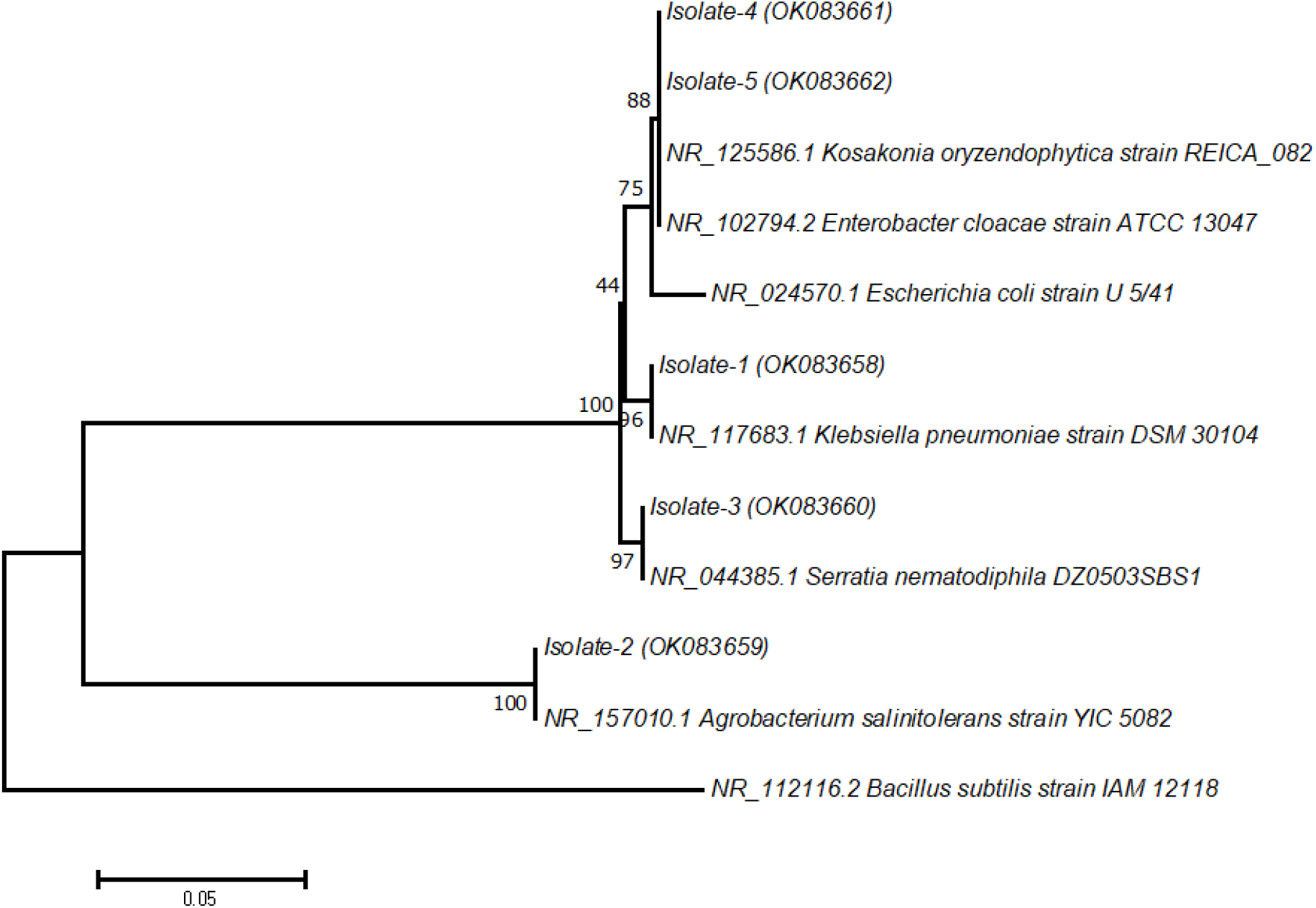
Maximum – likelihood phylogenetic tree showing the taxonomic placement and the evolutionary relationships of the five exoelectrogenic bacterial isolates from the MFC anodes isolated using the novel MnO_2_ based growth medium. Values at the nodes indicate the bootstrap support and the scale bar indicates the number of nucleotide substitutions per site.

Such Raman mapping experiments allow for accurate chemical characterization of biological material in their native state when they are contained within various biological structures such as biofilms, sludge flocs and tissues (Fernando et al., 2019, Huang et al., 2010 and Wang et al., 2016). Therefore, the outcomes of Raman analysis is direct evidence of electrochemically active bacteria over expressing vital cytochrome proteins when grown in the novel MnO_2_ containing medium or when forming a thick biofilm on the anode surfaces of functioning MFCs. On the contrary, the same analysis provided no evidence for the excessive presence of reduced cytochrome proteins in *E.coli* cells.

### 3.4. Molecular microbiological characterization of isolated exoelectrogenic bacteria from MFC anodes

Five bacterial species that grew and decolourised the novel MnO_2_ based growth medium were isolated and characterized using 16s rRNA molecular marker gene-based identification (Table-1). Their taxonomic assignments were done based on the 16s rRNA gene sequences contained within the NCBI GenBank database.

**Table - 1:** The bacterial isolates obtained using the novel MnO_2_ based solid medium from the MFC anode biofilms and their taxonomic assignments from 16s rRNA gene molecular microbiological analysis.

| MFC anode Isolate | NCBI accession number | Closest relative (Accession number of the closest match) | Percentage similarity (%) |
| --- | --- | --- | --- |
| Isolate-1 | OK083658 | *Klebsiella pneumoniae* strain DSM 30104 (NR_117683.1) | 100 |
| Isolate-2 | OK083659 | *Agrobacterium salinitolerans* strain YIC 5082 (NR_157010.1) | 99.86 |
| Isolate-3 | OK083660 | *Serratia nematodiphila* DZ0503SBS1 (NR_044385.1) | 99.75 |
| Isolate-4 | OK083661 | *Kosakonia oryzendophytica* strain REICA_082 (NR_125586.1) | 99.05 |
| Isolate-5 | OK083662 | *Enterobacter cloacae*. strain ATCC 13047 (NR_102794.2) | 99.74 |

Several environmental isolates of *Klebsiella pneumoniae* has previously been identified as electrochemically active and were demonstrated to be capable of conducting electron shuttling reactions within MFCs (Kim et al., 2019 and Zhang et al., 2008). The bacterium *Enterobacter cloacae* is also previously known to be an electrochemically active exoelectrogen that was capable of producing biogenic electricity in MFC systems (Rezaei et al., 2009 and Toczyłowska-Mamińska et al., 2015).

However, other isolated bacteria from MFC anodes using this novel MnO_2_ – based medium - *Agrobacterium salinitolerans, Serratia nematodiphila* and *Kosakonia oryzendophytica* were not previously known to be exoelectrogenic bacteria. Therefore, it can be concluded that the exoelectrogenic properties of aforesaid three bacterial species were hitherto unknown and have been conclusively demonstrated by the outcomes of this study.

### 3.5. Proposed name for the MnO_2_ – based exoelectrogen isolation/growth medium

The authors propose the usage of the name “E-Z exoelectrogen growth and isolation medium” for the newly developed microbiological solid medium for electrochemically active bacteria.

## 4. Conclusions

This study has demonstrated for the first time that electrochemically active exoelectrogenic bacteria can be conveniently isolated and grown using a MnO_2_ – based solid growth medium. Due the colour change of the medium from black to colourless when exoelectrogens reduce MnO_2_ in the medium, this medium offers the added benefit of a chromogenic growth/isolation microbiological medium. The outcomes of this work clearly demonstrated that exoelectrogenic bacteria such as *S.oneidensis* MR1 and other such environmental isolates can be clearly discriminated and distinguished from non-exoelectrogenic bacteria such as *E.coli* ATCC 25922. Further characterization of the MFC isolates by Raman micro spectroscopy conclusively demonstrated that the isolated bacteria using the novel medium were over expressing cytochrome proteins that are useful in extracellular electron transfer events.

## 5. Acknowledgements

EF and ZN would like to thank Professor G. R. A. Kumara of National Institute for Fundamental Studies (NIFS) for giving access to vibrational spectroscopy facilities throughout this study. Authors also would like to thank the Faculty of Applied Sciences, Rajarata University of Sri Lanka for providing material and logistical support for initiating and conducting this research.

